# Tissue-level integration overrides gradations of differentiating cell identity in beetle extraembryonic tissue

**DOI:** 10.1101/2024.04.10.588827

**Authors:** Katie E. Mann, Kristen A. Panfilio

## Abstract

During animal embryogenesis, one of the earliest specification events distinguishes extraembryonic (EE) from embryonic tissue fates: the serosa in the case of the insects. While it is well established that the homeodomain transcription factor Zen1 is the critical determinant of the serosa, subsequent realization of the tissue’s identity has not been investigated. Here, we examine serosal differentiation in the beetle *Tribolium castaneum* based on quantification of morphological and morphogenetic features, comparing embryos from a *Tc-zen1* RNAi dilution series, where complete knockdown results in amnion-only EE tissue identity. We assess features including cell density, tissue boundary morphology, and nuclear size as dynamic readouts for progressive tissue maturation. While some features exhibit an all-or-nothing outcome, other key features show dose-dependent phenotypic responses with trait-specific thresholds. Collectively, these findings provide nuance beyond the known status of Tc-Zen1 as a selector gene for serosal tissue patterning. Overall, our approach illustrates how analysis of tissue maturation dynamics from live imaging extends but also challenges interpretations based on gene expression data, refining our understanding of tissue identity and when it is achieved.

## 1. Introduction

Early patterning decisions form critical checkpoints in development. Changes in patterning can drastically alter tissue identity, structure, and function, as well as the chances of survival [1–4]. Furthermore, the progression from specification to tissue differentiation and maturation offers opportunities for the integration of multiple genetic inputs that can refine or alter upstream specification [5–7]. While certain developmental events require threshold amounts of key patterning genes [8], other aspects can proceed in a modular, independent fashion [9, 10]. Thus, investigating the phenotypic effects of patterning gene manipulation can distinguish mechanisms required for the full spectrum of tissue differentiation.

The differentiation of the extraembryonic (EE) tissues represents a key system for investigating patterning decisions during animal embryogenesis. These tissues are among the earliest to differentiate. Their rapid maturation is implicated in the evolutionary success of the amniote vertebrates and the insects, due to their essential roles in protecting and provisioning the embryo [11–13]. Specific properties of the fully differentiated tissues are tightly linked to their protective functions. For example, mouse syncytiotrophoblasts and the two insect EE tissues – amnion and serosa – switch from mitosis to the endocycle and become polyploid, with tissue-specific levels of ploidy [14]. Polyploidy in EE tissues is hypothesized to provide mechanical and physiological protection, as the large nuclei support tissue integrity and protection against infection, including via increased expression of signaling and effector molecules for innate immunity [14–17]. Additionally, the insect serosal tissue, as the outermost cellular layer in the egg, secretes a thick chitin-based cuticle that enhances eggshell strength and desiccation resistance [18–20].

In insects, the homeobox gene *zen* (or *zerknüllt*), the diverged orthologue of *Hox3*, is well established as a marker gene for early EE tissue [3, 21–23], and with clear evidence that it is the single critical EE determinant – a “selector gene” – in holometabolous species [1, 24]. It specifies the amnioserosa as the single EE tissue in cyclorrhaphan flies including *Drosophila melanogaster* [25, 26]. In fly species with a distinct serosa and a minimal amnion, Zen orthologues determine the early EE tissue region as serosal; subsequent, additional genetic inputs specify the amnion at the EE periphery and delimit the embryonic-extraembryonic tissue border [24, 27–29].

In the red flour beetle, *Tribolium castaneum*, the orthologue *Tc-zen1* is strictly required for serosal identity in an oblique anterior-dorsal region of the blastoderm (Fig. 1a-b; [1, 5, 30]). Subsequent morphogenesis leads to the mature EE tissue configuration: the outer serosa fully envelops the embryo, amnion, and yolk, while the inner amnion delimits a fluid-filled cavity ventral to the embryo (Fig. 1c; [31–33]). This configuration represents the standard EE tissue complement in winged insects [11, 34]. After *Tc-zen1* RNAi, loss of serosal identity leads to respecification of the blastoderm, with an anterior-dorsal expansion of amniotic and embryonic head tissue identities (Fig. 2a1,2e1; [1]).

**Fig. 1.**
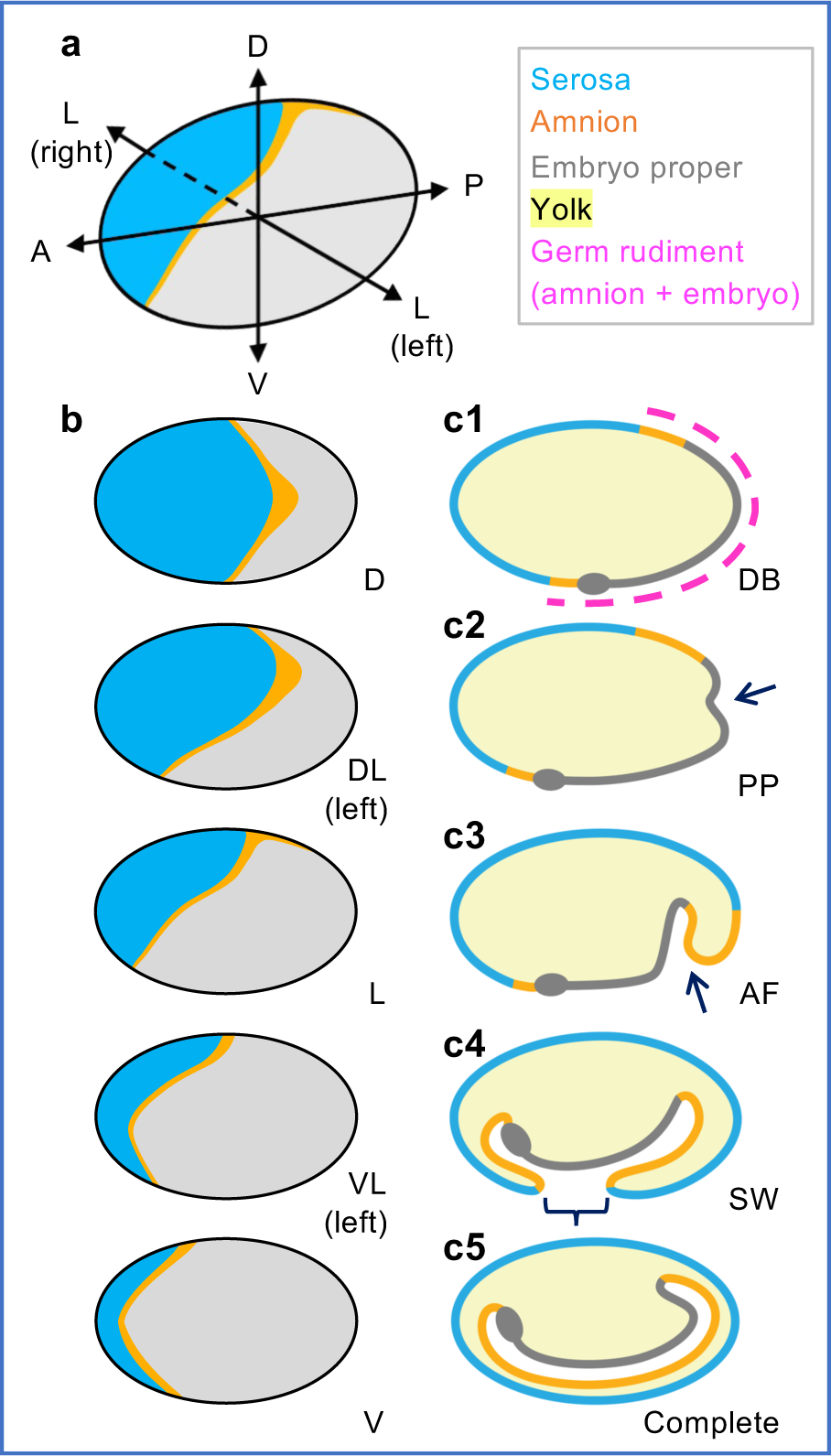
Early *Tribolium* wild type fate map and extraembryonic (EE) morphogenesis. **(a)** Axial orientation and **(b)** surface views at the differentiated blastoderm stage. The angle of view affects the visible area of the anterior-dorsal EE domain. The presumptive amniotic region is based on gene expression and lineage tracing [1, 43, 48]. **(c)** Ensuing morphogenesis leads to the embryo being fully enclosed in an outer serosa and inner amnion (mid-sagittal views): arrows indicate initial apical constriction and deeper invagination; curly bracket spans the open “window” region; grey ovals indicate the anterior of the embryo proper. Abbreviations: for angles of view: D, dorsal; DL, dorsal-lateral; L, lateral; VL, ventral-lateral; V, ventral; for landmark stages: DB, differentiated blastoderm; PP, primitive pit; AF, amniotic fold; SW, serosal window. Morphogenesis schematics modified from [13].

**Fig. 2.**
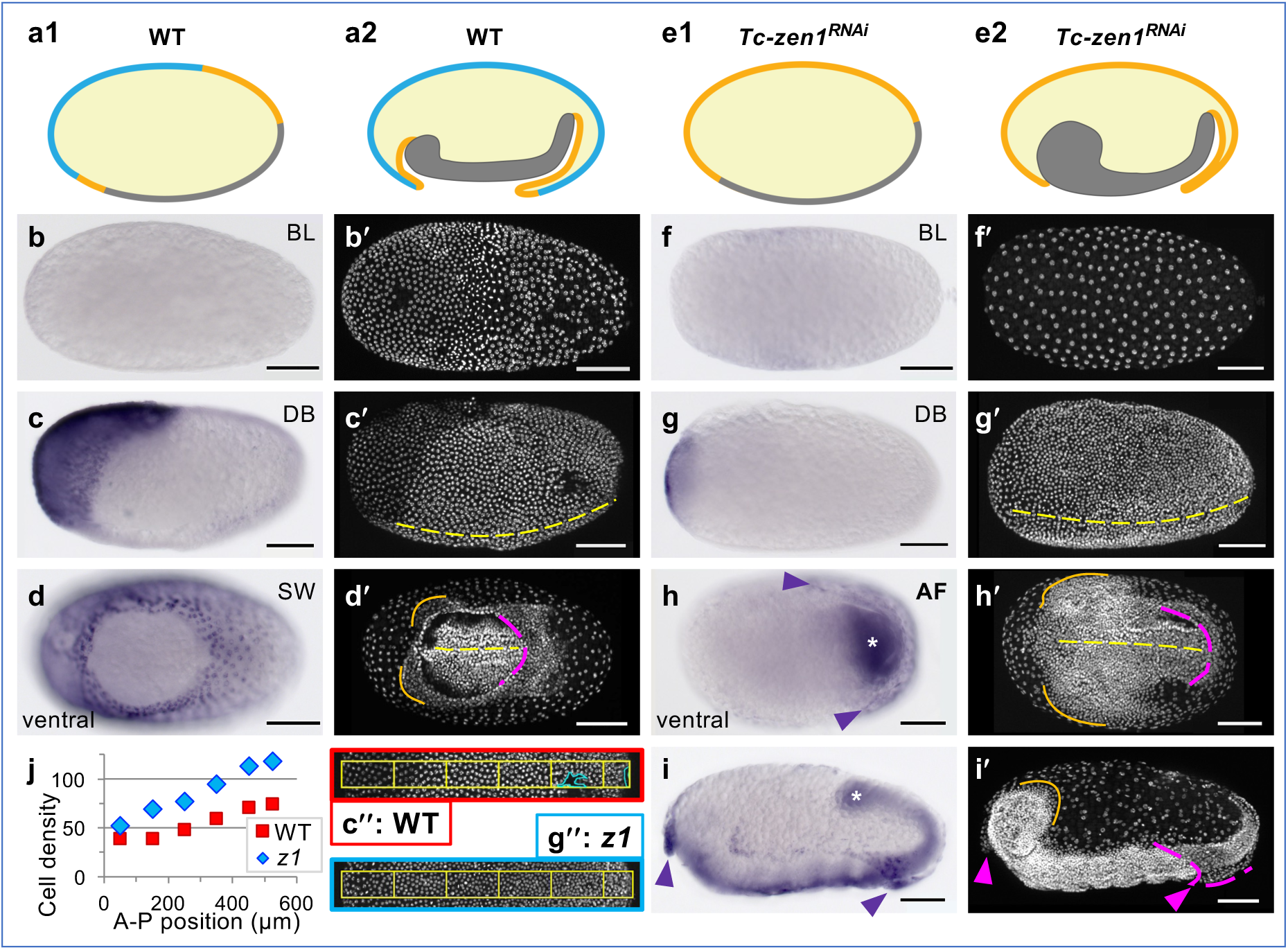
*Tc-hnt* expression reveals subtle tissue regionalization after *Tc-zen1* RNAi. Fate map, morphology, and expression of *Tc-hnt* in wild type (WT) **(a-d)** and *Tc-zen1* RNAi (“*z1*”) **(e-i)** embryos. **(a1-2, e1-2)** Mid-sagittal schematics of the serosal (blue), amniotic (orange), and embryonic (grey) domains at the differentiated blastoderm and extending germband stages. **(b-d, f-i)** *Tc-hnt in situ* hybridization, with DAPI counterstain (letter-prime panels): blastoderm formation **(b, f)**, differentiated blastoderm **(c, g)**, and EE tissue fold formation (**d:** closing serosal window, **h-i**: persistent ventral-posterior amniotic fold, “AF”). Views are ventral-lateral **(c, g)**, ventral **(d, h)**, lateral **(i),** or indeterminate **(b, f)**. Morphological annotations: <u>yellow:</u> ventral midline; <u>magenta</u>: posterior rim of the EE tissue enclosing the embryo; <u>orange</u>: head lobe rim; <u>arrowheads</u>: *Tc-hnt*-expressing folds of EE tissue after knockdown. Scale bars are 100 µm. **(j, c″, g″)**. Knockdown embryos exhibit higher cell density throughout the differentiating blastoderm and yet retain lower density in the ATR, which coincides with residual *Tc-hnt* expression (measured in 100 ξ 50 µm sectors along the anterior-posterior, A-P, axis, as depicted in c″ and g″ for the exemplar embryos shown in c-c′ and g-g′, respectively: double-prime images indicate measured sectors; density was scaled for sector size and to exclude minor damage: cyan regions).

Yet, the depth of characterization of EE tissue development in *Tribolium* has led to intriguing observations that belie the apparent all-or-nothing character of serosal specification by Tc-Zen1. First, Tc-Zen1 acts upstream or parallel to factors with dynamic expression waves that originate at the anterior pole and then pass posteriorly, exiting the presumptive serosal region and subsequently defining the adjacent presumptive amniotic tissue (*e.g.*, *Tc-Doc, Tc-iro*; [9, 35]). Second, *Tc-zen1* is zygotically expressed, downstream of a battery of maternal patterning factors for anterior and terminal system identities [2, 5, 36]. Thus, we wondered how modulating levels of *Tc-zen1* would impact serosal tissue specification against this backdrop of dynamic anterior gene expression.

Here, we assess hallmark serosal features throughout differentiation. Our marker gene profiling highlights an ostensible contradiction between early subregionalization and final tissue fate. To go beyond this, we then use a phenotypic series generated with a dilution series for *Tc-zen1* double-stranded RNA (dsRNA). Our live imaging approach to track active dynamics and quantify transient tissue features reveals subtle aspects of differentiation. We speculated that either the character of the presumptive serosal region shows progressive changes towards an amniotic identity with increasing dsRNA concentrations or that there is a distinct concentration threshold for serosal identity. In fact, we observe a series of thresholds for distinct aspects of differentiation, which can result in transient hybrid cell character, before the modular nature of EE identity progresses to a coherent, tissue-scale outcome.

## 2. Methods

### 2.1 *Tribolium castaneum* stocks

*Tribolium castaneum* (Herbst) beetles were maintained under standard conditions at 30 °C, 40-60% RH [37]. Strains used were San Bernardino (SB) wild type [37], nuclear GFP (nGFP) [38], and enhancer trap lines for the serosa (G12424, KT650), amnion (HC079), and cardioblasts/embryonic segmental domains (G04609) [33, 39, 40]. SB was used for *in situ* hybridization; the nGFP and enhancer trap lines were used for fluorescent imaging (Fig. 3).

**Fig. 3.**
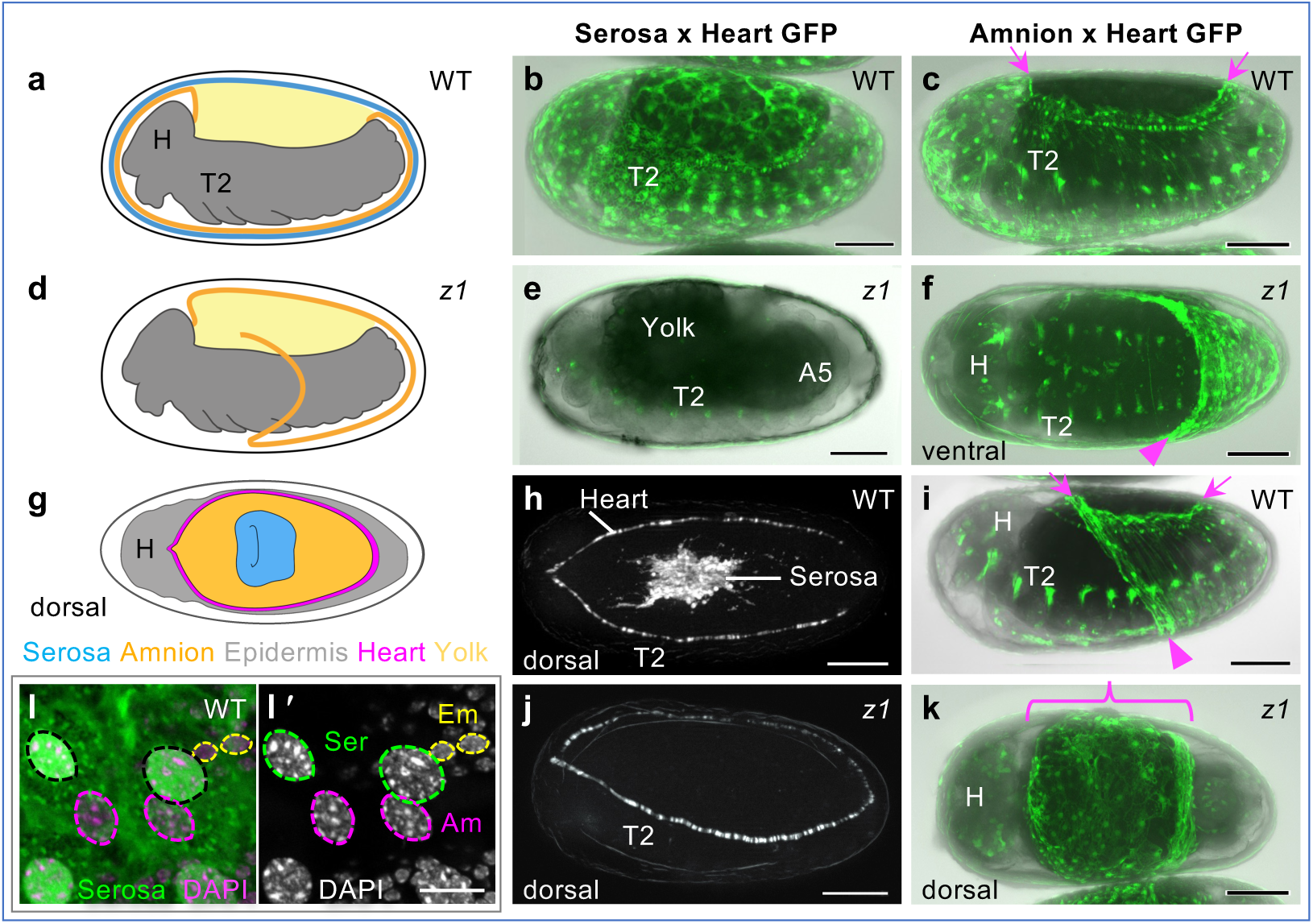
Strong *Tc-zen1* knockdown results in amnion identity in mature EE tissue. EE tissue structure and identity in WT **(a-c, g-i, l)** and after *Tc-zen1* RNAi **(d-f, j-k)**: (a-f, l) retracting and retracted germband stages, (g-i, j-k) EE tissue withdrawal and dorsal closure stages. Views are lateral unless otherwise indicated. In the schematic drawings, the outer-most dark grey layer is the eggshell (vitelline membrane). In the micrographs, GFP labels either EE tissue and a heart/segmental marker in heterozygous crosses, as indicated, in live (b-k) or fixed embryos (l). GFP signal overlaid on brightfield shows embryonic tissue as translucent grey and the yolk as opaque black. During dorsal closure, the cardioblasts delimit the EE region, obviating the need for brightfield (see also Movies 1-2). <u>Arrowheads</u> indicate amnion folded edges – of the persistent *Tc-zen1* RNAi ventral pouch (f) and transient WT ventral sac during mid-withdrawal (i). Before dorsal closure, the WT amnion does not cover the yolk (a, c, i: <u>arrows</u>), while knockdown EE tissue fully spans the dorsal region and is amnion-GFP-positive (d, k: <u>curly bracket</u>). Nuclear outlines are based on DAPI counterstain (l′); only serosal nuclei are GFP-positive (l). Abbreviations: Am, amnion; Em, embryo; H, head; Ser, serosa; T2, second thoracic segment. Scale bars are 100 µm (b-k) and 10 µm (l).

### 2.2 RNAi and *in situ* hybridization

Linear PCR amplicons served as template for dsRNA (Ambion T7 Megascript kit) and in *situ* hybridization probes (Roche DIG RNA Labeling Mix and T7 RNA Polymerase kits), as described [9, 33]. Sequences are based on the official gene set, OGS 3 [41]. Primer sequences are listed in Table S1, including adapters for adding T7 promotor sites [as in 33].

For RNAi, dsRNA for *Tc-zen1* (TC000921) used a 203-bp fragment for the eRNAi dsRNA dilution-series [as in 5, 42] and a 688-bp fragment for pRNAi in marker (trans)gene assays [as in 9, 39]. Experimental controls included uninjected adults or eggs (negative control) and RNAi for an unrelated gene that produced a wholly distinct knockdown phenotype (positive control: cell cycle regulator *Tc-double parked* (*Tc-dup*), TC003416, 373-bp amplicon; phenotype of severe blastoderm mitosis impairment, not shown). For eRNAi, dsRNA concentrations were: 100 ng/µl, 200 ng/µl, and 750 ng/µl for *Tc-zen1*; and 250 ng/µl and 1 µg/µl for *Tc-dup*, with consistent and minimal injection wound leakage, ensuring reasonable accuracy for dsRNA delivery. For pRNAi, *Tc-zen1* dsRNA was injected at 1 µg/µl. At least two biological replicates were performed for each RNAi treatment. For eRNAi, preblastoderm eggs (2-3 hAEL) were dechorionated in bleach (“DanKlorix Hygienereiniger”, Colgate-Palmolive: 4-5% sodium carbonate and 1-4% sodium hypochlorite), mounted in halocarbon oil 700 (Sigma), and injected anteriorly, as described [9]. For each eRNAi experiment, approximately 10 uninjected eggs and 20 eggs per RNAi sample condition were analyzed. For pRNAi, dsRNA was injected into the abdomen of adult females [20, 39].

For whole mount *in situ* hybridization, *Tc-hnt* (TC009560) was detected with a 627-bp probe [9]. Sense strand probe detection served as a negative control, while the previously characterized amniotic marker *Tc-pannier* was used as positive control (*Tc-pnr*, TC010407, [1, 43], 757-bp probe). Colorimetric detection was as described [9, 33]. Control eggs from uninjected mothers were stained in parallel with the *Tc-zen1* RNAi eggs, including matched duration of the alkaline phosphatase reaction for NBT-BCIP detection. All probes were used in at least three independent experiments. Eggs were imaged in Vectashield mountant with DAPI (Vector Laboratories), as described [5]. Focal stacking for *in situ* images was either done manually in Photoshop (Adobe) or with Helicon Focus (v.7.5.8, Helicon Soft).

### 2.3 Live imaging acquisition and analysis

Time-lapse movies were acquired with a Zeiss AxioImager.Z2 with Apotome.2 structured illumination epifluorescent microscope, as described [20], at 26-28 °C in intervals of 15 or 30 minutes. Data handling was performed in Image J (NIH), including cell tracking with the MTrackJ plugin [44]. Unscorable eggs (*e.g*., unfertilized or non-specific defects) were excluded from subsequent analyses. Maximum intensity z-projected images were used for all analyses. Eggs were categorized by angle of view to ensure consistent comparisons. Sample sizes and numerical results are specified in the figure legends and Supplementary File 1.

#### 2.3a Spatial and temporal analyses

Early embryos were manually labeled at the boundary between the non-dividing EE tissue with low cell density and the condensing germ rudiment tissue with high cell density (Fig. 4). Traces of the boundaries were then overlayed to illustrate the curvature and variation in border shape per treatment (Fig. 4i-p). For sufficient sampling, embryos that varied between strictly dorsal and dorsal-lateral angles of view were included, which is reflected in minor skewing in the traces. EE tissue area was normalized to egg area (μm^2^) from the z-projected images. Aspect ratios were based on maximum axial extent of the EE tissue relative to total egg length (anterior-posterior, A-P) and width (dorsal-ventral, D-V).

**Fig. 4.**
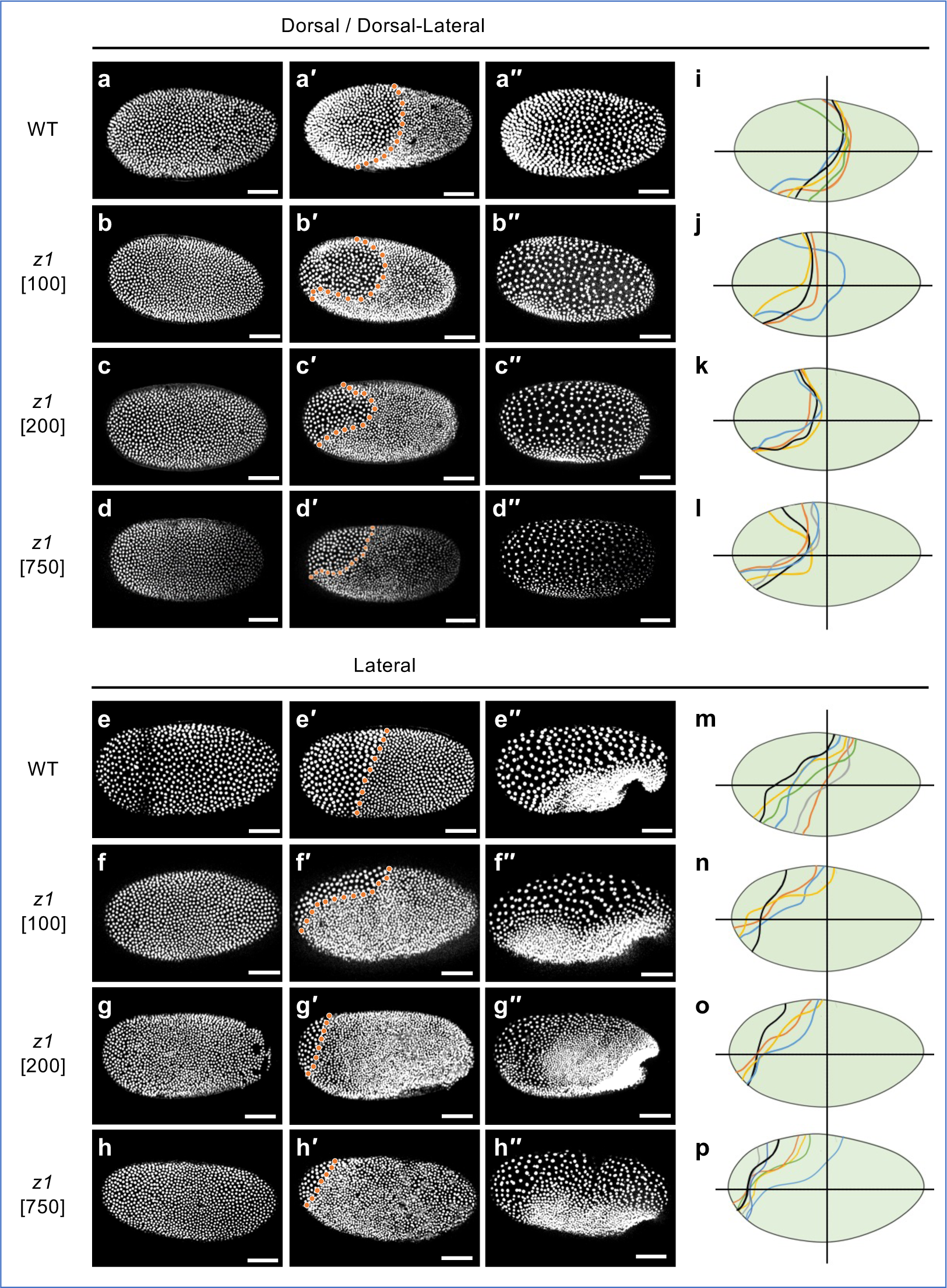
*Tc-zen1* dsRNA concentration-dependent reduction in the morphologically distinct EE domain. Qualitative analysis of EE tissue area at the uniform blastoderm **(a-h),** differentiated blastoderm **(a′-h′)** and the posterior fold stages **(a′′-h′′).** Orange dotted line marks the boundary between the EE and condensing embryonic tissues. EE tissue area, in both dorsal (a-d) and lateral (e-h) aspect, is greatest in WT (uninjected controls) and progressively reduces across the three dsRNA treatments (concentrations as indicated: 100, 200, or 750 ng/µl; see also Movies 3-7). Although there is no evident subregionalization within the WT EE tissue (presumptive serosa), the EE tissue in strong knockdown embryos (presumptive amnion, with 750 ng/µl) is reduced to the small, persistent ATR. **(i-p)** EE domain traces depicting boundary shape and curvature for all treatments at the differentiated blastoderm stage: each colored line represents one embryo in the dataset (see Methods for boundary determination). Horizontal lines indicate the A-P axis midline; vertical lines at 50% egg length provide a landmark for comparing EE tissue extent. Scale bars are 100 μm.

The timing of blastoderm differentiation was when the germ rudiment-specific mitotic division occurs. If the differentiation occurred over several time frames of the movie, the median frame was taken. The formation of the posterior amniotic fold was determined as the earliest time frame when the invaginating primitive pit had deepened sufficiently to form a ventrally oriented tissue crest that could be consistently discerned from multiple angles of view (as in Fig. 4g′′). Movie time frames were converted into age (hours after egg lay: hAEL) to directly compare treatments, based on minimum age from a 1-hour egg collection.

#### 2.3b Nuclear area and cell density

To measure nuclear size (apical area on z-projected images), contrast settings were adjusted for consistency of GFP appearance across embryos, and nuclei were manually outlined with the polygon tool in ImageJ, excluding nuclei at the periphery to avoid skewing due to egg curvature. For establishing baseline differences between EE and germ rudiment tissue, 3 nuclei were measured for both tissues in 6-16 embryos per treatment condition. For measuring EE tissue alone at selected stages, 10 nuclei were measured per embryo from the same dataset. To ensure balanced sampling across the EE tissue, at the blastoderm differentiation stage nuclei were randomly chosen within each of four equally sized subregions (illustrated in Fig. S1). At the extended germband stage (20 hAEL), nuclei covering the yolk were selected from throughout the tissue. Lastly, for tracking changes in EE nuclear size over time, 5 nuclei were tracked for 3 embryos per treatment condition.

Cell (nuclear) density in the mature EE tissue (20 hAEL) was determined with the multi-point tool in Image J, within three square regions (80 µm x 80 µm; illustrated in Fig. S2). Counts included nuclei for which >50% of the nuclear area was within the bounding box. For statistical analysis, the mean count from the three regions per embryo was used.

#### 2.3c Head lobe area and cell density in the EE-embryonic tissue border region

Morphology was quantified for head size and EE rim cell packing – the cell density transition from embryonic to EE tissue in the region of the uncovered head lobes. Head area (µm^2^) was measured at 18-20 hAEL (at the extended germband stage, morphologically matched by eye within this stage range/time) for one head lobe fully visible in lateral aspect. The head region was delimited as follows: the dorsal surface was taken as the maximum region within which cell density was sufficiently high that individual nuclei could not be distinguished. The posterior extent of the head was delimited by the inflection point in shape between the head lobe and the rest of the germband. The ventral extent was measured as follows: to account for minor differences in angle, the region of interest for the head area was measured either to the ventral edge of the egg or to the morphologically distinct ventral midline, whichever was the smaller region.

To systematically sample EE cell density radiating out from the condensed head tissue of the embryo (illustrated in Fig. 7a**′**-b**′**, below), five rectangular regions of interest (85 x 16 µm) were equally spaced, radiating in step sizes of 25°. Rectangles were positioned adjacent and tangential to the dense embryonic tissue region, defined as the region within which nuclear density was too great to distinguish individual nuclei. Nuclei were counted as described above (including nuclei with ≥50% of total nuclear area inside the region of interest).

### 2.4 Statistical analysis

Shapiro Wilk tests determined the normality of the data. For non-normally distributed data that required >2 comparisons, the Kruskal-Wallis test (non-parametric one-way ANOVA) was used to compare distribution of the data, with Dunn’s post hoc analysis for pairwise comparisons of means using rank sums. The Bonferroni correction was used to adjust the significance values for multiple comparisons. The Mann-Whitney U test was used for data from two independent samples. For normally distributed data that required >2 comparisons, the one-way ANOVA tested for variation in the means, with post hoc Tukey HSD (honestly significant difference) for pairwise significance. Independent sample *t*-tests were used for comparisons between two treatment groups only. Statistical tests were conducted in GraphPad Prism (version 9.5.0) and graphs were made in Microsoft Excel. In the figures, significance levels are: *: p<0.05, **: p<0.01, ***: p<0.001. Box plots depict the mean (“X”), median (horizontal line), and Q1-Q3 interquartile range (box height); bars and whiskers denote the full data range, with outliers plotted individually. All values for graphs and statistical test details are provided in Supplementary File 1.

## 3. Results

### 3.1 Early morphogenetic and morphological differentiation of the serosa

Early embryogenesis has been well characterized in *Tribolium* [1, 5, 9, 31, 32, 42, 45]. Briefly, formation of the <u>uniform blastoderm</u> as a continuous epithelium over the yolk culminates in completion of cellularization and optimization of cell packing (Fig. 2b,b′; [46, 47]). Then, the 13^th^ cell division results in the <u>differentiated blastoderm</u>, distinguishing the germ rudiment, which remains mitotically active, from the serosa, which ceases mitosis and enters the endocycle (Fig. 1c1). In parallel, early morphogenesis begins. This involves rapid condensation of the embryo and its invagination into the yolk while serosal epiboly maintains tissue continuity over the yolk surface. Key morphogenetic features of this process are: (1) formation of the <u>primitive pit (PP)</u> via apical constriction at the posterior pole, initiating invagination, (2) the anteriorly advancing ventral <u>amniotic fold (AF)</u> that engulfs the embryo, and (3) final closure of the <u>serosal window (SW)</u> opening, utilizing an EE supracellular actomyosin cable and including contributions from minor anterior and lateral EE folds (Fig. 1c2-1c4). Severing their physical connection, the serosa and amnion then form a double membrane surrounding the embryo proper, producing the mature EE tissue topology (Figs. 1c5, 2a2) by 12 hAEL (17% of embryogenesis, [33]).

Morphologically, blastoderm differentiation rapidly results in a visual distinction between the serosa and germ rudiment (Fig. 2c,c′,c″). As morphogenesis proceeds, serosal cells flatten and become squamous, with low cell density and large, polyploid nuclei – defining features of the serosa (Fig. 2d,d′,j; [31, 32]). After modest further mitosis during germband extension, the amnion also enters the endocycle, although amniotic nuclei remain smaller, with characteristically lower ploidy than the serosa (Fig. 3, below; [20, 39, 48, 49]).

### 3.2 Strong *Tc-zen1* RNAi leads to loss of serosal tissue identity

After *Tc-zen1* RNAi, the blastoderm anterior is respecified, and early morphogenesis results in partial enclosure of the embryo by folds of the amnion alone (Fig. 2e1-e2; [1, 20]). Here we reproduce this full strength *Tc-zen1* RNAi knockdown phenotype (Fig. 2f-i), which has been described previously from pupal [1, 5], adult [9, 20], and embryonic [42] injections.

Key morphological hallmarks of this strong phenotype are germ rudiment-specific mitotic divisions occurring throughout an expanded region of the blastoderm (Fig. 2g,g′,g′′,j) and characteristic head lobe and EE tissue folding morphology. Specifically, the dorsal-lateral rim of the head lobes exhibits diffuse cell packing (Fig. 2h,h′) and remains uncovered by EE tissue folds (Fig. 2i,i′). In contrast, the EE tissue forms an extensive ventral fold, or pouch, over the thorax and abdomen, and there is often a medial fold, or flap, of EE tissue that overhangs the head at the anterior pole (Fig. 2h-i).

We further confirm that the EE tissue in the strong *Tc-zen1* knockdown is amnion. This was previously shown with the early amniotic marker genes *Tc-pannier* (*Tc-pnr*) [1] and *Tc-Dorsocross* (*Tc-Doc*) [9], which expand throughout the EE tissue domain in *Tc-zen1* RNAi embryos. However, these genes are pleiotropic and later acquire embryonic expression domains. To determine the identity of the mature EE tissue after *Tc-zen1* RNAi, we conducted a live imaging screen with transgenic lines with tissue-specific GFP (Fig. 3). We visualized either the serosa or the amnion in heterozygous crosses with an embryonic enhancer trap line that provided an imaging internal control, labeling the heart (cardioblasts) and body segments as anatomical landmarks (Fig. 3a-c). We examined embryos throughout the middle 50% of embryogenesis (24-60 hAEL), from the extended germband through EE withdrawal and dorsal closure stages (Fig. 3a,d,g; Movie 1). With two independent serosa-GFP lines, after *Tc-zen1* RNAi we observe a complete absence of GFP in EE tissue (100%, n= 170, Fig. 3b,e,h,j; Movie 2). In contrast, amnion-GFP is expressed throughout the knockdown EE tissue, including in the early EE ventral pouch and the entire dorsal region during EE withdrawal (n= 47, Fig. 3c,f,i,k). In the mature EE tissues, wild type serosal nuclei are two-thirds larger than amniotic nuclei (apical area), and this distinction persists after knockdown (Fig. 3b,c,k,l,l′; [20]). Lastly, we detect no serosal cuticle (mechanical reinforcement of the eggshell) after *Tc-zen1* RNAi in any of the transgenic backgrounds [as in 5, 19, 20].

Thus, tissue-specific transgenes as well as nuclear size and absence of cuticle confirm the stable identity of all EE cells as amniotic after strong *Tc-zen1* RNAi, supporting the mid-embryogenesis morphological hallmarks as reliable indicators for this phenotype.

### 3.3 *Tc-zen1* RNAi reveals early, subtle regionalization within the EE tissue

Despite the clear amniotic identity of *Tc-zen1* RNAi EE tissue, based on multiple genetic and morphological features, the serosal marker gene *Tc-hindsight* (*Tc-hnt*) reveals regionalization within the knockdown tissue that challenges a binary serosa/amnion tissue identity. Wild type expression of *Tc-hnt* arises in the serosa during blastoderm differentiation and persists as a faithful marker of the entire tissue throughout germband stages (Fig. 2b-d, [9, 35]). After strong knockdown of *Tc-zen1* and loss of serosal identity, *Tc-hnt* is largely absent but still detected in two specific domains (Fig. 2f-i). First, *Tc-hnt* has restricted expression in morphogenetically active folds of the EE tissue (Fig. 2h-i: arrowheads), consistent with regulation of ectopic supracellular actin cables by *Tc-Dorsocross* [9].

More intriguingly, after *Tc-zen1* RNAi, early *Tc-hnt* expression still occurs in a small, transient domain at the anterior pole (Fig. 2g). This domain may reflect *Tc-zen1*-independent patterning [4, 9, 36]. Yet, this genetic distinction correlates with morphology that is characteristic of the serosa: this region has larger cell nuclei (Fig. 2g,g**′**) and notably lower cell density than the rest of the blastoderm (Fig. 2c′′,g′′, j: anteriormost region; [1]). Thus, we wondered how varying the strength of *Tc-zen1* RNAi would alter this distinctive anterior terminal region (ATR) and resulting tissue specification.

### 3.4 A phenotypic spectrum for EE tissue differentiation

To look at spatiotemporal dynamics with accuracy of stage-matched comparisons between wild type and knockdown embryos, and to move beyond marker gene analyses restricted to specific candidate (trans)genes (Figs. 2-3), we next undertook a quantitative live imaging approach to examine the EE tissue in a dilution series for *Tc-zen1* dsRNA. Our primary focus was on morphological and morphogenetic features that define tissue identity, and how this is manifest particularly in the ATR. These experiments were conducted in a transgenic line in which ubiquitously expressed, nuclear-localized GFP fully labels nuclear volume [38]. This allowed us to visualize all nuclei (all cells) of the blastoderm, irrespective of differentiation status or subsequent tissue identity, and to quantify key diagnostic features such as timing of mitosis, cell packing, and nuclear size (Movie 3).

We first asked whether it is possible to achieve weaker knockdown phenotypes for *Tc-zen1*, and if so, what the consequences would be for blastoderm differentiation and early tissue morphogenesis. To explore this, we performed embryonic RNAi (eRNAi) by injecting three distinct concentrations: 100, 200, or 750 ng/µl, balancing the need for statistically robust sample sizes with testing a range of dsRNA dilutions. With our highest eRNAi concentration, representing 75% of the dsRNA concentration previously used in both eRNAi [42] and parental RNAi (pRNAi: *e.g.*, Fig. 2), we could reproduce the full-strength phenotype (Fig. 4, and discussed below). Our lower concentrations represent five-fold and ten-fold dilutions compared to previous eRNAi full-strength *Tc-zen1* knockdown [42]. These concentrations did indeed produce a range of outcomes for EE tissue area, which is most evident during blastoderm differentiation (Fig. 4a′-h′,i-p; Movies 4-7).

To ensure that our quantitative analyses of EE structure are directly comparable, we evaluated the timing of early developmental events. eRNAi for *Tc-zen1* does not significantly affect the timing of blastoderm differentiation with any of the three dsRNA concentrations (Fig. 5a). As the wild type controls were uninjected, this also provides evidence that the eRNAi treatment itself did not alter the rate of early development. In contrast, as the *Tc-zen1* RNAi phenotype manifests and develops, this led to a significant delay in EE tissue folding in the strongest knockdown treatment by 17% development (Fig. 5b: an 80-minute delay compared to wild type, p=0.012, Mann Whitney U test; see Supplementary File 1 for all raw data values and test statistic details here and below).

**Fig. 5.**
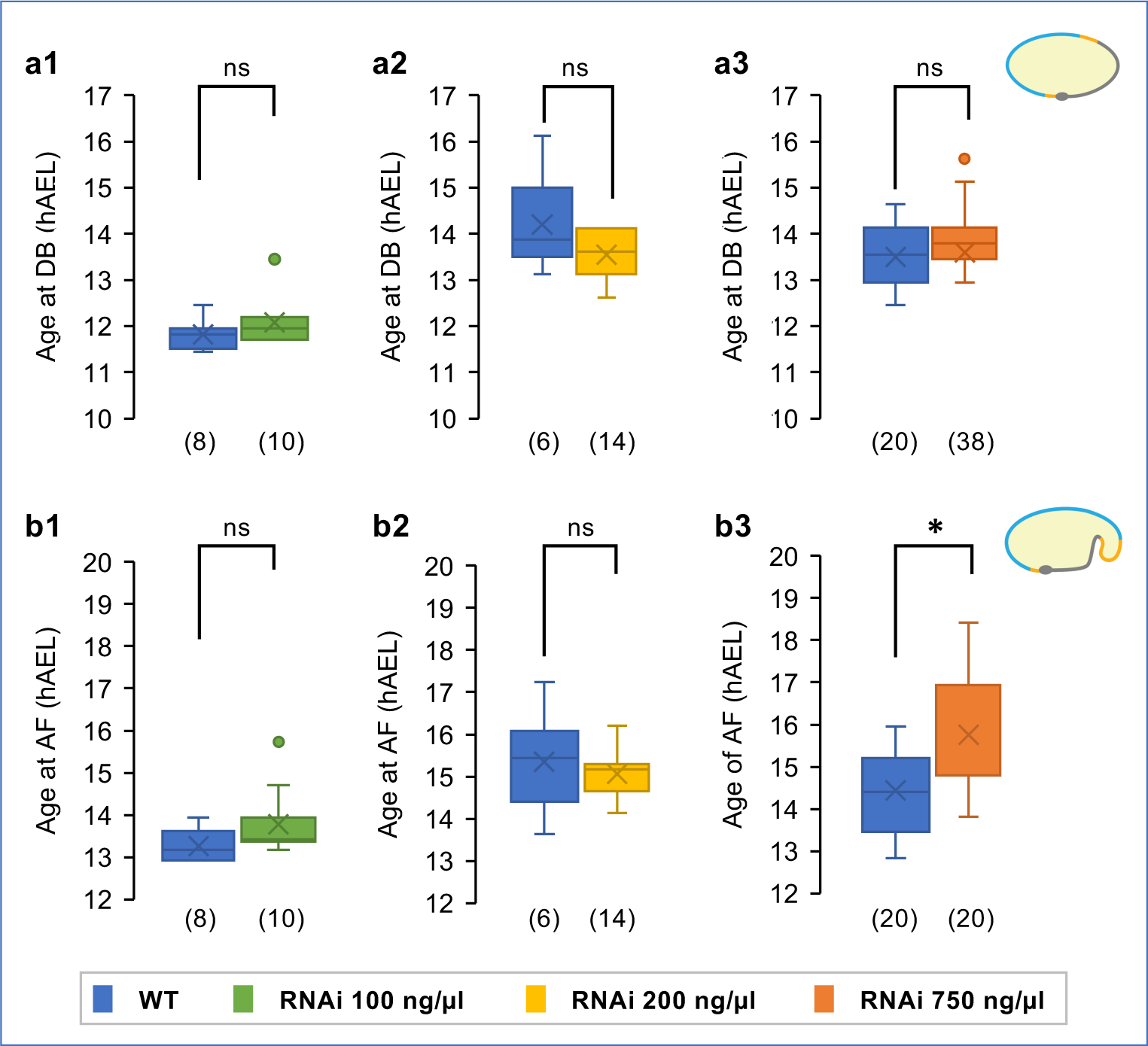
Timing of landmark early events. Timing of blastoderm differentiation (**a1-a3**: DB) and amniotic folding (**b1-b3**: AF). Only embryos in strictly lateral view were analyzed for the latter. To mitigate interexperimental differences such as ambient temperature, knockdown embryos were directly compared to within-experiment WT controls. **S**ignificance was determined by Mann-Whitney U tests, revealing the onset of a delay during amniotic fold formation in the strongest knockdown treatment (p<0.05). Sample sizes are indicated parenthetically; box plot features are defined in the Methods. See Supplementary File 1 for raw values and test statistics.

### 3.5 Dose-dependent and non-uniform reduction in the EE domain after *Tc-zen1* RNAi

To examine EE tissue size and shape, we first inspected embryos during early differentiation (Fig. 4a-h**′′**). Wild type embryos (Fig. 4a**′**,e**′**) have a large serosal domain with low cell density and large nuclei, whereas strong *Tc-zen1* knockdown retains only the small ATR of presumptive amniotic tissue (Fig. 4d**′**,h**′**). Embryos injected with the lower dsRNA concentrations display EE area of intermediate sizes (Fig. 4b-b**′′**,c-c**′′**,f-f**′′**,g-g**′′**). We noted natural, inter-embryo variability in EE tissue area, but did not observe any systematic left-right asymmetries or treatment-specific changes in boundary curvature (Fig. 4i-p).

Going further, we observe an overall graded decrease in EE tissue area with increasing concentrations of *Tc-zen1* dsRNA, with threshold sensitivities depending on embryo aspect (angle of view: Fig. 6a). The wild type serosa covers half the dorsal surface area and one-fifth ventrally (Figs. 1, 6a). The strongest knockdown treatment (750 ng/µl) resulted in significant reduction in EE tissue area for all five angles of view from dorsal to ventral (p≤0.019, non-parametric Kruskal-Wallis with Dunn’s post hoc test). Although mean EE tissue area was visibly reduced with both lower dsRNA concentrations (Figs. 4i-p, 6a), reduction was only statistically significant for the 200-ng/µl treatment, in most but not all angles of view (p≤0.047), in part due to statistical power when finely parsing our dataset sample sizes by angle of view. Despite this limitation, we were intrigued that in dorsal aspect we observed a consistent 40% reduction in mean EE tissue area compared to wild type with all knockdown treatments (range: 36-44%), whereas reduction in ventral aspect ranged over two-fold (40%-76%), in a dsRNA dose-dependent manner. Furthermore, the 200-ng/µl treatment reveals differing thresholds for degree of EE tissue reduction, with values comparable to the 100-ng/µl treatment (dorsal and dorsal-lateral), comparable to the 750-ng/µl treatment (lateral and ventral-lateral), or intermediate (ventral view).

**Fig. 6.**
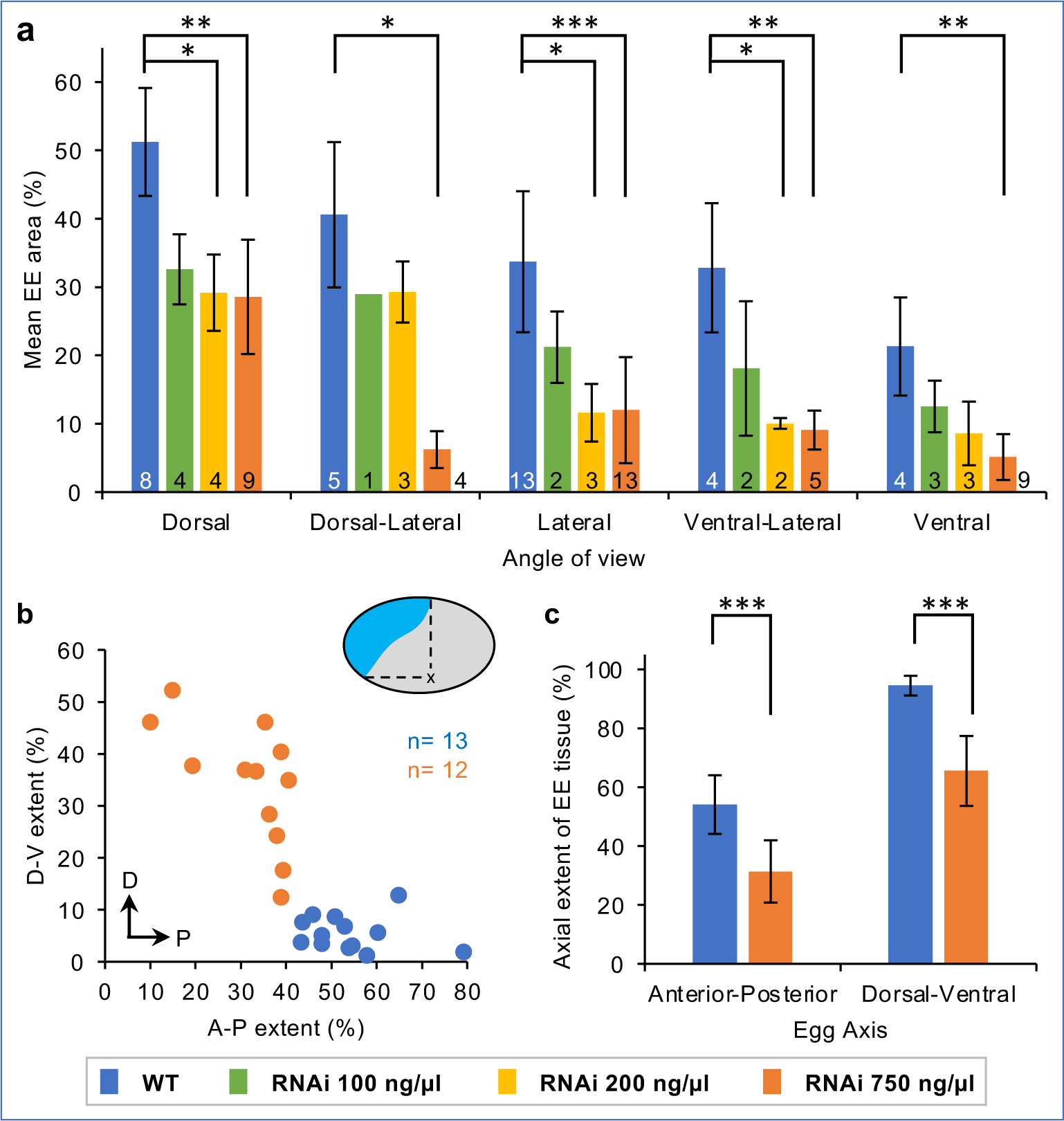
Non-uniform and threshold responses in knockdown EE area reduction. **(a)** Mean EE tissue area (% relative to egg size) at the differentiated blastoderm stage, for all angles of view for each RNAi condition, with sample sizes indicated at the base of the graph. Error bars represent ± one standard deviation. **(b)** An aspect ratio scatter plot for individual embryos in lateral aspect shows relative reduction of EE tissue area along the A-P and D-V axes (sample sizes indicated in the plot). Compared to the WT serosa, after strong knock-down (dsRNA 750 ng/μl) the presumptive EE tissue does not extend as far ventrally or posteriorly. **(c)** Mean ± standard deviation summary statistics for the dataset in panel (b). Significance levels are defined in the Methods; unlabeled pairwise comparisons were not significant. See Supplementary File 1 for raw values and test statistics (a: Kruskal-Wallis with Dunn’s post hoc; c: Mann-Whitney U).

To evaluate the non-uniform reduction in EE tissue after *Tc-zen1* RNAi, we also assessed EE domain size along both the A-P and D-V axes (Fig. 6b). The wild type serosa spans half the egg length dorsally and nearly the entire egg width in the anterior (Fig. 6b-c: mean of 54% A-P and 94% D-V, compare with Fig. 4a**′**,e**′)**. In contrast, strong *Tc-zen1* RNAi embryos have a significantly reduced EE tissue area along both axes, but with a greater absolute reduction along the D-V axis (Fig. 6b-c: mean of 31% A-P, and 66% D-V; p<0.0001 for both axes: Mann-Whitney U tests).

Altogether, this indicates a dose-dependent effect of *Tc-zen1* knockdown on specification and resulting EE area, with non-uniform changes in the position of the EE-embryo tissue boundary.

### 3.6 Head morphology shows conflicting trends for knockdown phenotypic response

If we have achieved weaker knockdown phenotypes with lower concentrations of *Tc-zen1* dsRNA, what are the implications for serosal *vs.* amniotic identity of the EE tissue? As EE tissue structure resulting from complete loss of serosal identity is highly characteristic at germband stages (Figs. 2-3), we directly quantified these morphological hallmarks in our dilution series dataset (Fig. 7a-b**′**).

**Fig. 7.**
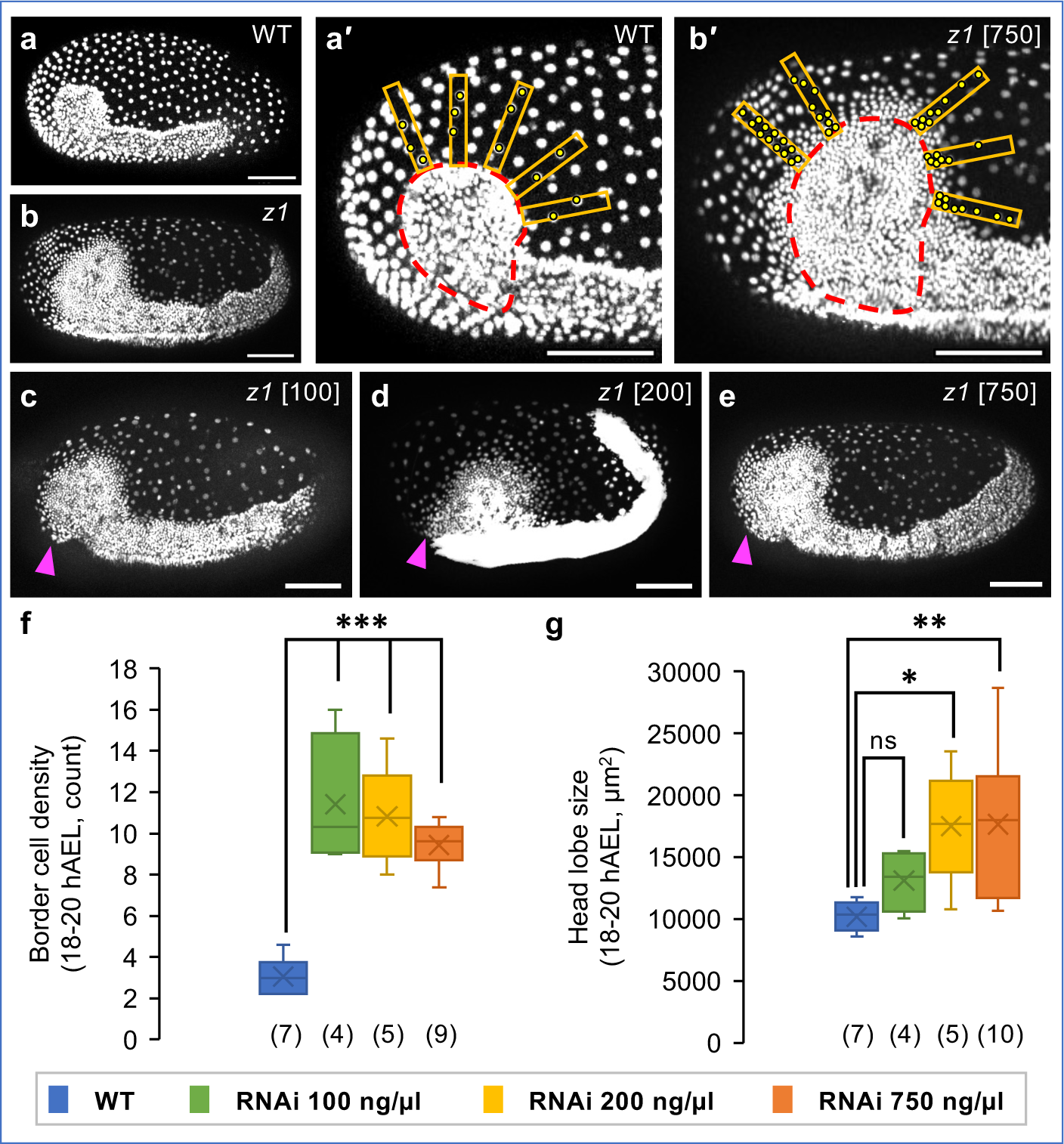
Evaluation of head size and peripheral cell packing. **(a-b)** Lateral aspect at 19 hAEL for WT (a) and strong *Tc-zen1* knockdown (b), depicting the extended germband embryo and widely spaced EE tissue. **(a′-b′)** Images annotated for quantification of head size (dashed red outline) and cell packing (density) at the EE-embryonic border (orange rectangles with yellow nuclear counts: see Methods). **(c-e)** Stage-matched knockdown embryos (dsRNA concentrations as indicated), illustrating head size, head rim cell density, and the anterior-medial EE tissue flap (arrowheads). All scale bars are 100 µm. **(f-g)** Quantification of head rim cell density (f) and head area (g: single head lobe). Sample sizes are indicated parenthetically; box plot features and significance levels are defined in the Methods. All RNAi treatments were significantly different from WT but not from one another for border cell density. See Supplementary File 1 for raw values and test statistics (f-g: One-way ANOVA and post-hoc Tukey’s HSD).

With all three dsRNA concentrations, we find that knockdown embryos have (1) enlarged heads that (2) remain uncovered by EE tissue except in an anterior-medial flap and with (3) diffuse cell packing along the dorsal-lateral rim of the head lobes (Fig. 7a-g).

For head rim cell packing, all three knockdown treatments were comparable to one another, with a significant, >3-fold increase in border region cell density compared to wild type embryos (Fig. 7f). By this stage, in fact the wild type embryo is fully enclosed by and physically separate from the outer serosa, while after *Tc-zen1* RNAi the head lobe ectoderm and EE tissue are contiguous within a cell sheet on the egg surface. Nonetheless, our quantification strategy addresses how compact, or diffuse, the embryonic tissue rim is, as well as local cell density within the anterior EE tissue. Curiously, among the knockdown treatments, we see a counter-intuitive trend of a minor increase in head lobe rim delineation (lower cell density due to a sharper tissue boundary transition) and with less inter-embryo variability with increasing concentrations of *Tc-zen1* dsRNA. Despite care in stage-matching embryos, these non-significant differences may reflect subtle differences in developmental rate for head tissue condensation [1]. Alternatively, an intriguing possibility is that this may reflect delays due to greater ambiguity of tissue identity in the weaker knockdown conditions.

As with differentiated EE tissue area (Fig. 6a), germband-stage head size reveals a graded response to knockdown, and again with the 200-ng/µl treatment more similar to the strong knockdown treatment (Fig. 7g). The weakest knockdown treatment shows a modest 1.3-fold increase in head area compared to wild type. Both the 200-ng/µl and 750-ng/µl treatments resulted in a more substantial, and significant, 1.7-fold increase in head size.

Taken together, the features of head rim cell packing and absence of a lateral EE cover over the head lobes strongly distinguish all *Tc-zen1* knockdown conditions from the wild type. This suggests that the knockdown embryos feature a complete loss of serosal identity and that the EE territory comprises of strictly amniotic tissue. However, the phenotypic outcome for head size supports a graded, dose-dependent knockdown response.

### 3.7 *Tc-zen1* RNAi influences the rate of EE polyploidization

Finally, we examined nuclear size dynamics as a fine-scale diagnostic feature of EE tissue identity, as nuclear size scales with ploidy under stage- and tissue-matched conditions (Fig. 3l; [31, 50]).

During wild type blastoderm differentiation (about 13 hAEL), germ rudiment nuclei divide and become less than half the size of the non-dividing serosal nuclei (Fig. 8a-a′,b). All knockdown treatments also show a significant, >2-fold nuclear size difference between the ATR and condensing embryonic tissue (Fig. 8c-e). EE nuclei in all knockdown treatments were about 8% smaller than wild type serosal nuclei, although this early difference was not significant (Fig. 8f: wild type mean of 61 µm^2^; knockdown EE means of 54-58 µm^2^; see Fig S1 for method). This confirms observations after strong knockdown (*e.g*., Fig. 2g**′**), with the retention of larger nuclei in the ATR even with complete loss of serosal identity.

**Fig. 8.**
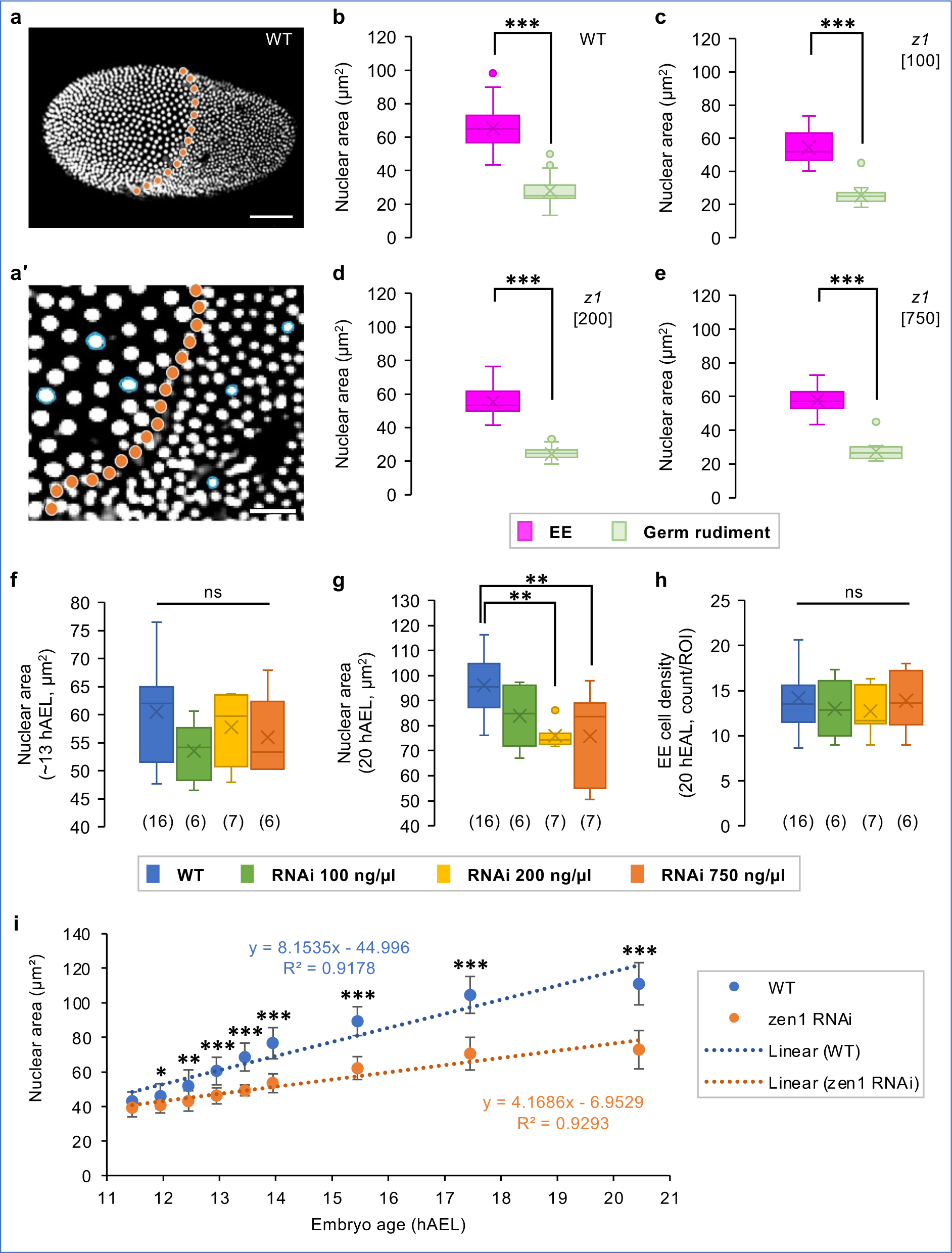
EE nuclear size dynamics as a proxy for tissue-specific polyploidization. **(a-a′)** At blastoderm differentiation, WT serosal nuclei (anterior, left) are larger than germ rudiment nuclei (posterior, right). The orange dotted line marks the tissue boundary; blue circles delineate selected nuclei (inset image; see Methods). Scale bars are 100 µm (a) and 25 µm (a′). **(b-e)** Nuclear area in the EE tissue and germ rudiment at the differentiated blastoderm stage (treatments indicated in the figure panels), with 3 nuclei measured per tissue region in 6-16 embryos per treatment (sample sizes as specified in panel f). **(f-g)** EE nuclear area at the differentiated blastoderm (13 hAEL) and extended germband (20 hAEL) stages, with 10 EE nuclei measured per embryo (sample sizes are indicated parenthetically; method schematic in Fig. S1). **(h)** EE tissue cell density at 20 hAEL, as mean nuclear count for three EE regions per embryo (each region 80 µm x 80 µm: method schematic in Fig. S2). **(i)** EE nuclear size over time. For each treatment, 5 EE nuclei were tracked in 3 embryos (n=15 nuclei/treatment), plotted as mean ± standard deviation (see Fig. S4 for individual plots). Equations for the linear trendlines and correlation coefficients (R^2^) are shown. See Supplementary File 1 for raw values and test statistics (b-e: Mann Whitney U; f-h: one-way ANOVA; i: *t*-test). Box plot features and significance levels are defined in the Methods.

However, by the time of maximum germband extension (20 hAEL), we do detect significant differences in EE nuclear size (Fig. 8g). Consistent with entry into the endocycle, EE nuclei increased in size in all casaes, but to a lesser extent in the 200-ng/µl and 750-ng/µl treatments. In the wild type and 100-ng/µl treatments, EE nuclei increased 1.6-fold, albeit with the RNAi sample still proportionately smaller. In contrast, in the two stronger knockdown treatments, EE nuclei increased by only 1.3-fold, attaining only 80% of wild type serosal nuclear size (76 µm^2^ *cf*. 96 µm^2^). Despite this binary difference in EE nuclear size increase, cell density within the EE tissue does not differ across treatments (Figs. 8h, S2).

Given the different EE nuclear sizes between control and low dsRNA compared to the medium and high dsRNA treatments by the stage of germband extension (Figs. 8g, S3), we analyzed the dynamics of nuclear size increase for each of these groups (Figs. 8i, S4). From 12 hAEL onward, wild type serosal nuclei are significantly larger than the 750-ng/µl EE nuclei. Moreover, there is a 2.0-fold faster rate of nuclear size increase in wild type throughout germband extension, heightened to 2.4-fold in a fast phase immediately after blastoderm differentiation (Supplementary File 1). In contrast, knockdown EE nuclei exhibit a modest, gradual increase – consistent with amnion-like nuclei that are ultimately smaller and with weaker GFP fluorescent signal than the serosa (Figs. 8i, S3; [20]).

In sum, our live imaging quantification suggests that wild type EE nuclei enter the endocycle at a faster cycle rate compared to after strong *Tc-zen1* knockdown. As EE ploidy levels are tissue-specific, this dynamic feature argues that weak knockdown EE tissue is hybrid in character between bona fide serosa and amnion – exhibiting serosal-like nuclear size increase (Fig. 8g) in a tissue that is structurally amnion-like (Fig. 7c,f, 8h).

## 4. Discussion

Morphological appearances of cells and their genetic identities are normally coherent. For example, mammalian immune cells can be clearly distinguished by their nuclear morphology, degree of granulation, and relative size [51]. Furthermore, cell morphology can define functional state and predict subsequent fate, such as in pre-cancerous cells [52, 53].

In this study, we use the insect EE epithelia as a tractable research system to explore (a) the limits of marker gene expression for assessing tissue identity after fate map changes in developing tissues and (b) how weaker RNAi phenotypes can reveal the effect of modulating a selector gene transcription factor on tissue identity and structure.

### 4.1 Uncoupled and dose-dependent features of tissue identity

Genetic markers revealed complete loss of serosal tissue identity after *Tc-zen1* RNAi (Figs. 2-3; [1, 5]), but with two unexpected outcomes. First, the blastoderm ATR is genetically but not morphologically distinct from a serosal identity (Fig. 2g,j). Second, altered gene expression in later development may reflect *functional* transdifferentiation – not of overall tissue identity, but of morphogenetic character – between serosa and amnion (Fig. 2i; [9]).

Our live imaging of a knockdown dilution series directly tackles these ambiguities, revealing subtle properties of tissue patterning and specification dynamics. Similar to analysis of dsRNA concentration thresholds for embryonic phenotypes [54], our treatment conditions dissect nuances of EE tissue specification. The earliest features – timing of blastoderm differentiation and EE nuclear size at differentiation (Figs. 5a, 8b-f) – are not predictive diagnostic features. However, both early EE and later embryonic tissue area support a graded phenotypic series after *Tc-zen1* knockdown (Figs. 4, 6a, 7g). It is striking that these different measures of tissue area are concordant, considering the difference between the two-dimensional simplicity of the blastoderm and later, multilayered and folded tissue geometries. Similarly, nuclear size dynamics reveal a threshold between the 100-ng/µl and 200-ng/µl treatments for whether nuclei are serosa-like or amnion-like in size and rate – but not onset – of polyploidization (Fig. 8g,i). And, only the 750-ng/µl treatment showed a slowing in EE morphogenesis (Fig. 5b3). This latter aspect likely reflects early secondary consequences of the largest shift to amniotic identity: amnion cells have a slower rate of cell shape change during embryonic gastrulation, presumably in part due to loss of anterior *fog* expression after loss of serosal identity [1, 31, 45]. Thus, a phenotypic series ranging from serosal to amniotic EE identity is supported by both cellular and tissue-scale properties, with specific tissue features sensitive to different levels of *Tc-zen1* knockdown.

Yet, mature tissue topology and structure are consistent with the complete loss of serosal identity in all knockdown treatments (Figs. 7c-f). Upon reflection, this suggests a fundamental difference in nuclear size regulation between the amnion and serosa. Nuclei that are more amnion-like are smaller (Figs. 3l, 8g,i, S3, S4), but neither EE cell density near the head lobes nor throughout the tissue distinguishes between knockdown treatments (Figs. 7f, 8h). This is despite the fact that nuclear:cell scaling would be expected to favor a higher density for cells with smaller nuclei [55]. This suggests that a smaller nuclear size (lower ploidy) is a refractory, inherent feature of the amnion that does not respond to changes in cell packing and that can be uncoupled from other EE tissue features. Also, the folds of knockdown EE tissue (Figs. 2i, 7c-e) may provide flexible reservoirs that ensure the dorsal yolk is fully covered by EE tissue, regardless of starting cell numbers.

Despite the knockdown ATR (Fig. 2g,j), there is no equivalent subregionalization in the wild type serosa (Fig. S3; [5, 9, 35]). Furthermore, even with transitional EE tissue sizes after knockdown (Figs. 4, 6), we still observed a clear EE-embryo tissue boundary. Also, nuclear size across the EE tissue exhibited no gradient that might imply mixed or progressive serosal/amniotic identities (Figs. 4 a**′**-h′, 8b-e, S3). A substantially weaker knockdown might produce transitional EE phenotypes. However, our observed phenotypes likely reflect binary EE/germ rudiment fate determination, integrating maternal genetic inputs.

### 4.2 Maternal terminal inputs pattern anterior tissue domains

As in *Drosophila*, the anterior pole of the *Tribolium* blastoderm is patterned by components of the terminal system. Maternal *Tc-torso-like* (*tsl*) activates zygotic *Tc-zen1, Tc-homeobrain* (*hbn*), and *Tc-decapentaplegic* (*dpp*) [2, 36]. Additionally, an anterior maternal gradient of the conserved Wnt inhibitor *Tc-axin* suppresses posterior, embryonic Wnt signaling and thereby promotes anterior fates [2, 4]. *Tc-hbn* and *Tc-zen1* then form a zygotic feedback loop, further suppressing posterior *Tc-caudal* (*cad*) and specifying the serosa [2]. The requirement of the terminal system to trigger this patterning cascade is further supported by expansion of the serosal domain after loss of the Torso pathway inhibitor *Tc-capicua* (*cic*) and its reduction after loss of the Torso downstream target *Tc-maelstrom* (*mael*) [56, 57]. Interestingly, while these factors affect the size of the serosal domain, their loss of function does not eliminate the serosa-germ rudiment tissue boundary, and none except *Tc-zen1* directly dictates serosal identity.

The majority of Torso pathway target genes remain to be identified [57]. One possible candidate is *Tc-hnt*, which retains ATR expression after *Tc-zen1* RNAi (Fig. 2g). This was an unexpected result, as wild type EE expression of *Tc-hnt* is exclusively serosal [9, 35]. However, comparison across species suggests that *hnt* expression dynamics involve *zen*-independent EE regulatory inputs for activation and spatial restriction. In *Drosophila*, amnioserosal expression of *Dm-hnt* does not fully coincide with *Dm-zen* but only occurs in a nested subdomain [58]. In the jewel wasp *Nasonia vitripennis*, earliest *Nv-hnt* expression is extensive and dynamic, before refining to the serosa in a domain that coincides with *Nv-zen* [59]. In the mosquito *Anopheles gambiae*, *Ag-zen* is strictly serosal whereas early *Ag-hnt* expression spans the presumptive serosal and amniotic domains [27]. In the scuttle fly *Megaselia abdita*, *Ma-hnt* also spans the early serosal and amniotic domains [60]. Although EE *hnt* expression begins early in all species, functional studies have mainly linked its function to later morphogenesis, but not patterning, in *Tribolium* and fly species, in addition to conserved late roles in the peripheral nervous system [9, 58, 60].

Together, maternal factors initially pattern anterior terminal gene expression and morphology, and the extent of this is unmasked after knockdown of *Tc-zen1*. However, in light of subsequent amnion EE identity after *Tc-zen1* knockdown, the anterior-terminal distinction represents a transient feature that is insufficient to dictate subsequent tissue identity.

### 4.3 A selector gene at the nexus of maternal patterning and zygotic EE specification

The selector gene concept distinguishes transcription factors that are necessary and sufficient to confer specific cell and tissue identity. They can override the “ground plan” genetic landscape of target cells and act in parallel on cells with differing ground states, to pattern a complex tissue in a unified and functionally integrated manner [61, 62]. For example, in murine neuronal stem cells, homeodomain selector genes act as global regulators across a complex regulatory landscape in their target cell types [63].

Classic examples of selector genes are the Hox genes that confer segment-specific identity along the anterior-posterior body axis of the embryo. The diverged insect *Hox3* orthologue, *zen*, or *zen1*, no longer functions in the embryo, and yet these genes retain conserved Hox-like sequence features and are recognized as having selector gene function in the extraembryonic domain [24, 34, 64, 65]. Hence, despite the complexity of maternal anterior patterning through multiple signaling pathways and other transcription factors, zygotic Zen alone is known to determine serosal tissue identity. In the fly *Megaselia*, ectopic *Ma-zen* promotes serosal development, in part through repression of amnion marker genes that respond to different thresholds of *Ma-zen*, and in *Drosophila* prolonged overexpression of *Dm-zen* leads to enlarged nuclei in the amnioserosa [60].

Similarly, in *Tribolium* we show here that while hallmark serosal cell morphology may originate independently of *Tc-zen1* activity in the ATR, this alone is insufficient to produce bona fide serosal identity (Figs. 2-3), and neither does this ground state alter mature tissue structure within the wild type serosa (Fig. S3). Thus, full serosal identity overrides the initial distinction of the ATR subregion. On the other hand, although the hybrid cell character in our weakest knockdown treatment exhibited serosal-like nuclear size increase (Figs. 8, S3), ultimately the tissue has amnion-like properties akin to the strong knockdown condition, including for integrated measures of nuclear morphology (Figs. 7, S3c).

In light of serosal tissue biology and function [5, 16, 39], we have uncovered different suites of downstream serosal target genes with different sensitivities to Tc-Zen1 activity. Whereas features such as cuticle secretion require high levels of Tc-Zen1, the initial nuclear size increase – but not full extent of polyploidization – responds to very low doses. As in a recent study of *Drosophila* limb patterning [66], we thus find that the importance of the selector gene as a master regulator is modulated by interplay with its wider genetic network, to ultimately pattern the full spectrum of tissue identity.

## STATEMENTS and DECLARATIONS

## Supporting information

Supplementary File 1. Raw data values and statistical tests from Figures 5-8.

Movie 1: WT dorsal closure

Movie 2: zen1 dorsal closure

Movie 3: WT lateral

Movie 4: z1_750 lateral

Movie 5: z1_200 lateral

Movie 6: z1_100 lateral

Movie 7: z1_100 dorsal

## Acknowledgments

We thank Thorsten Horn for guidance in performing the eRNAi injections; and Matthias Pechmann, Tim Saunders, and Siegfried Roth for discussions and feedback on the manuscript.

## Data availability

All data generated or analyzed during this study are included in this published article (and its supplementary information files).

## Author contributions

KEM devised statistical approaches, analyzed the data, and wrote the paper. KAP conceived of the study, generated and analyzed the data, and wrote the paper.

## Funding

This work was supported by funding from the Biotechnology and Biological Sciences Research Council (BBSRC, UKRI) through grant BB/V002392/1 and by core funding from the University of Hohenheim to KAP.

## Conflict of interest

The authors declare no competing interests.

## Supporting Information

### Additional supplementary files

Supplementary File 1. Raw data values and statistical tests from Figures 5-8.

Movie 1. Time-lapse movie of wild type dorsal closure in a heterozygous cross for serosal and cardioblast GFP, as in Fig. 3h. Posterior embryonic fluorescence includes the proctodeum (also visible in Fig. 3k), which flexes ventrally out of the field of view as dorsal closure progresses.

Movie 2. Time-lapse movie of dorsal closure after *Tc-zen1* parental RNAi, in a heterozygous cross for serosal and cardioblast GFP, as in Fig. 3j. There is minor fluorescent signal of yolk components, but a complete absence of serosal GFP. Without the contractile force of the serosa, dorsal closure is slower and less efficient than in wild type [20].

Movies 3-7. Representative time-lapse movies of early embryogenesis in the ubiquitous nuclear-GFP background for wild type (Movie 3) and each of the three *Tc-zen1* dsRNA concentrations (Movie 4: 750 ng/µl; Movie 5: 200 ng/µl; Movies 6-7: 100 ng/µl), from blastoderm differentiation through germband extension, as in Figs. 4 and 7. Shown in lateral aspect (Movies 3-6) or dorsal-lateral aspect (Movie 7). Note that fixed brightness settings on these export files leads to some overexposure of embryonic tissue in later stages, whereas in the original high resolution files brightness is within the acquisition dynamic range for visualizing the EE nuclei at all stages and for embryonic tissue at early stages (*e.g.*, Fig. S4b). The loss of peripheral signal in the last few hours of Movie 6 was due to a drop in focal plane of acquisition, not a loss of tissue in the embryo. Time stamps indicate minimum age from a 1-hour range, with recordings made at 26-28 °C.

**Fig. S1.**
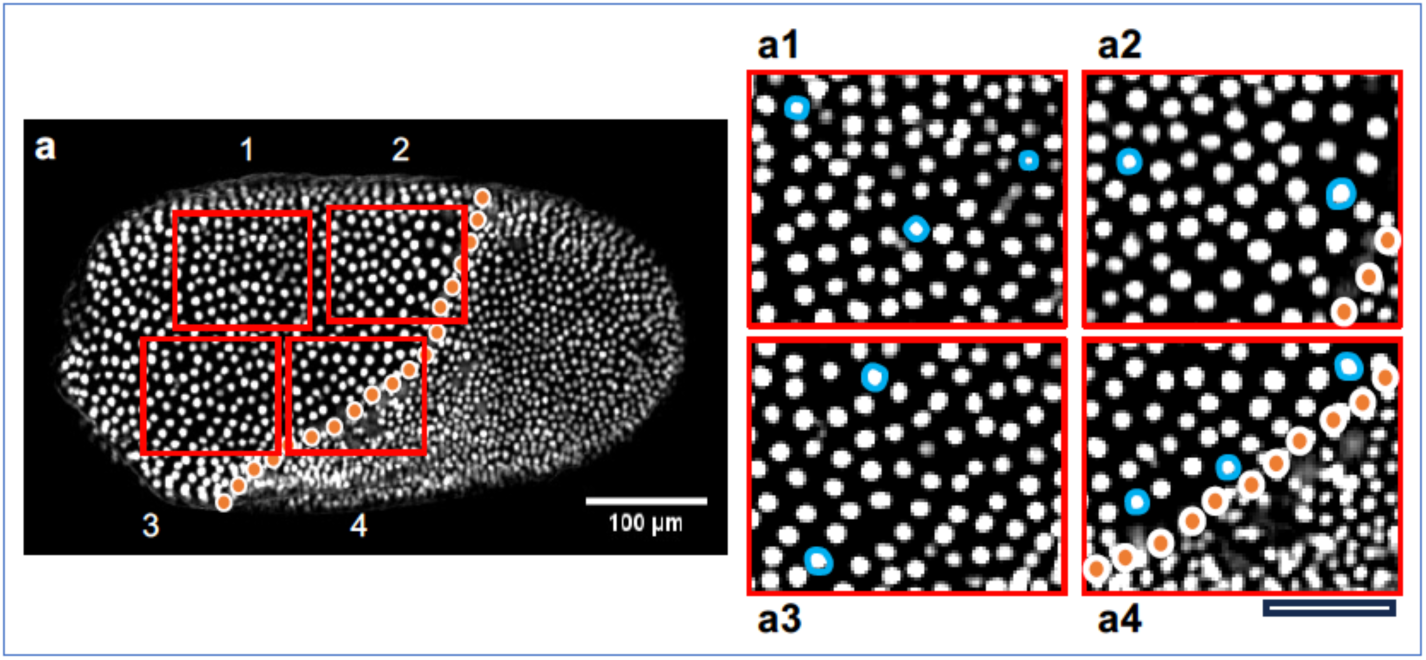
Measuring EE nuclear area during blastoderm differentiation. For each embryo, four rectangular regions (125 µm x 100 µm) were placed on the EE tissue territory and a total of 10 nuclei (2-3 nuclei per region) were chosen at random for quantification (inset images a1-a4), illustrated here with a wild type control embryo. At the extended germband stage (20 hAEL), nuclei covering the yolk were selected. Scale bar for all inset images is 50 µm. Supports Fig. 8f-g.

**Fig. S2.**
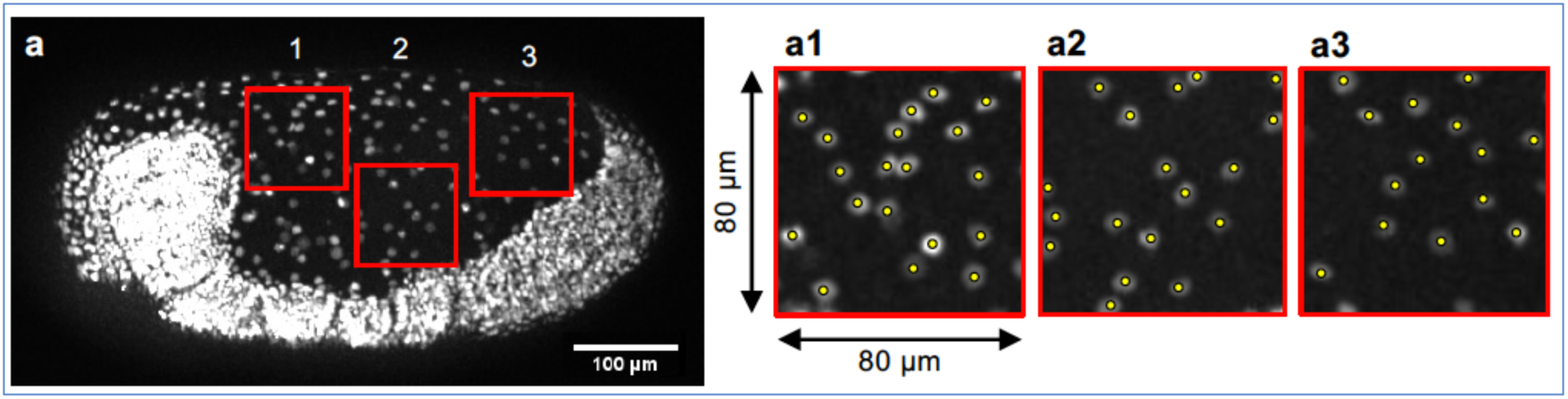
Measuring cell density of the EE tissue at 20 hAEL. For each embryo, three square regions (80 µm x 80 µm) were placed on the EE tissue territory and individual nuclei were counted, for all nuclei with >50% of nuclear area within the bounding boxes (inset images a1-a3), illustrated here with an embryo from the strong *Tc-zen1* RNAi treatment (750 ng/µl). The mean nuclear count from the three regions was recorded as the EE cell density value per embryo. Supports Fig. 8h.

**Fig. S3.**
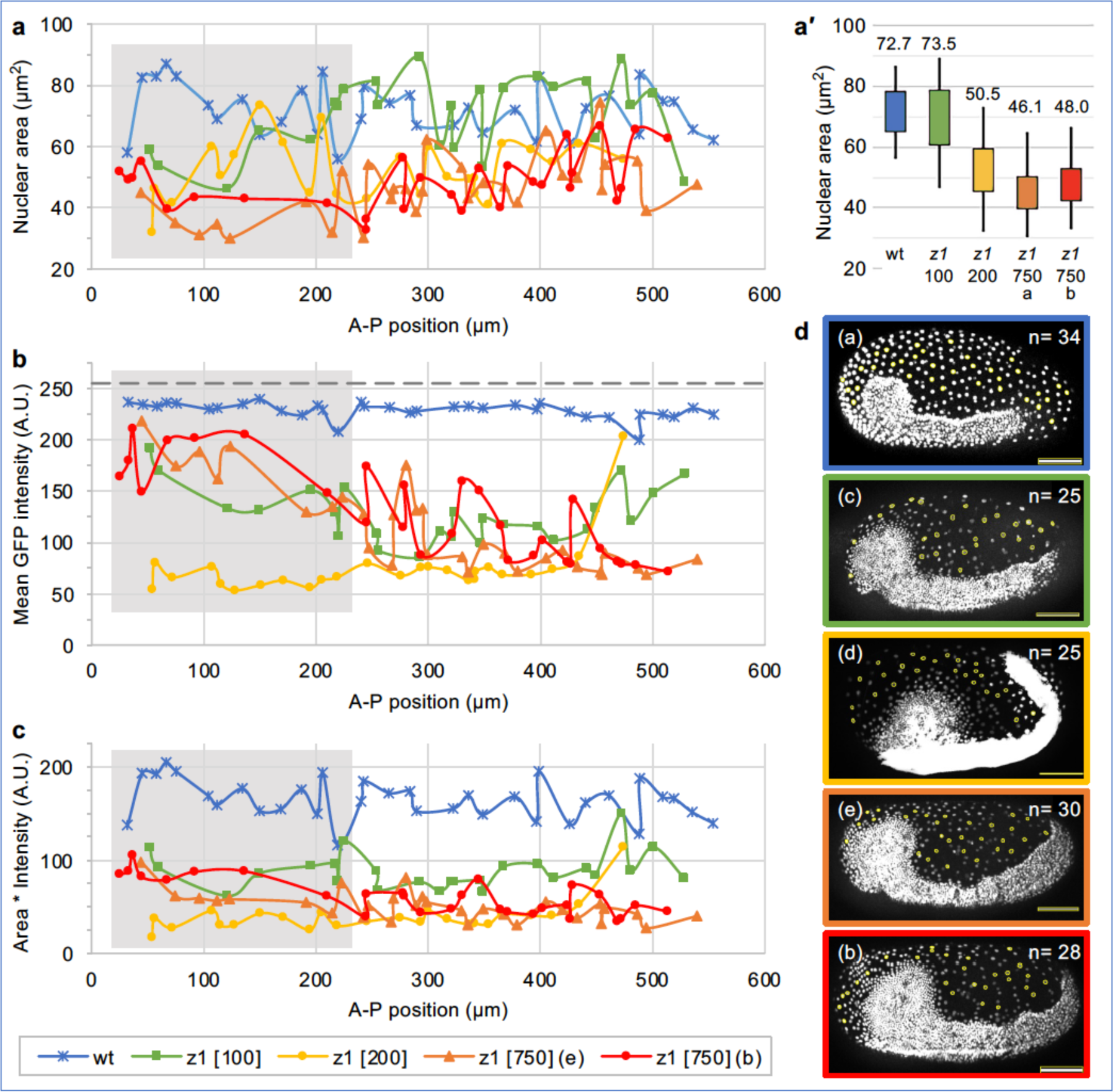
EE nuclear features mapped along the A-P axis at 19 hAEL. For each embryo, ≥25 EE nuclei were measured for size (**a**: area, µm^2^), mean GFP intensity (**b**: grayscale range of 0-255, with saturation at 255 indicated by the dashed black line), and for area weighted for intensity (**c**), based on the representative embryos for each treatment that are featured in main text Fig. 7 (**d**: corresponding main text image panel and nuclear sample sizes are indicated, with yellow circles for the measured nuclei). Additionally, for nuclear area the box plot (**a′**) depicts the interquartile range and minimum and maximum individual values, with the median reported within the chart. In (a-c), the grey shaded region indicates the position of the embryonic head lobes. Note that brightness settings are optimized for consistent, clear signal in the border region at the rim of the head lobes, leading to ostensible oversaturation in thick, multinuclear germband embryonic tissue, particularly as knock-down EE nuclei generally have weak fluorescence. Knockdown EE nuclei near embryonic tissue (nuclei measured in the most anterior and posterior regions) tend to be brighter (b), but not to differ in size compared to EE nuclei elsewhere in the tissue (a). Overall, nuclear area alone distinguishes between knockdown treatments, with the weakest knockdown comparable to wild type serosal nuclear size (a, a′), as in Fig. 8g; GFP intensity sharply distinguishes wild type serosal nuclei by brightness and homogeneity compared to all knockdown EE nuclei (b); while area weighted by GFP intensity most consistently distinguishes phenotypic categories for wild type, the weakest knock-down (100 ng/µl dsRNA), and stronger knockdown (200 or 750 ng/µl dsRNA) treatments. Supports Fig. 8f-g.

**Fig. S4.**
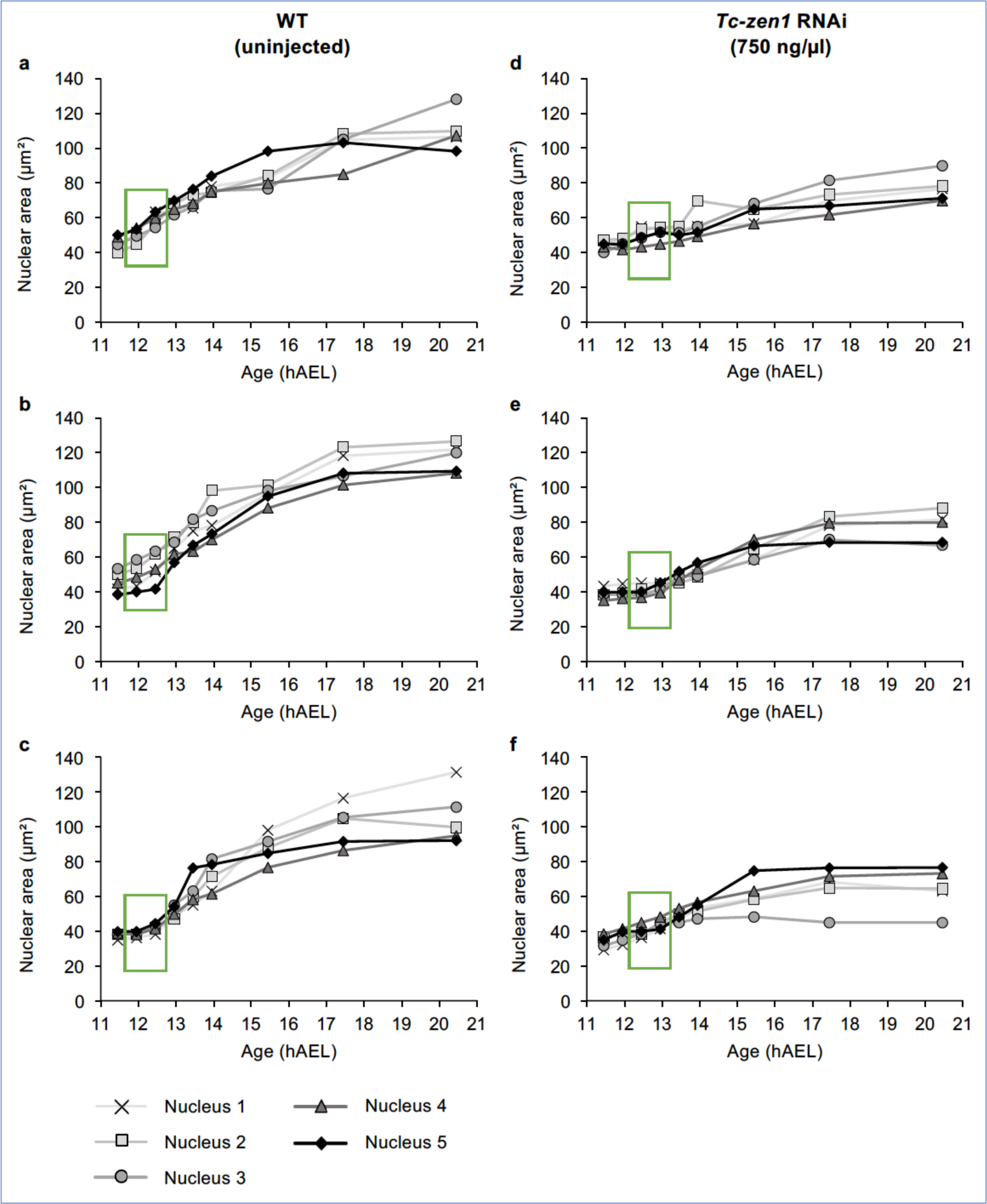
Nuclear area over time for WT and strong *Tc-zen1* knockdown embryos. Individual values for changes over time in nuclear area, for five tracked EE nuclei per embryo, in each of three embryos for the WT **(a-c)** and *Tc-zen1* RNAi 750 ng/µl **(d-e)** treatment conditions. Nuclei were tracked from 11.45 hAEL (prior to differentiation of the blastoderm) to 20.45 hAEL (stage of maximum germband extension), at nine selected time points. The green boxes demarcate the exact timing of blastoderm differentiation in each embryo. These data support Fig. 8i, where the average of the 15 nuclei per treatment is plotted with the standard deviation.

**Table S1.**
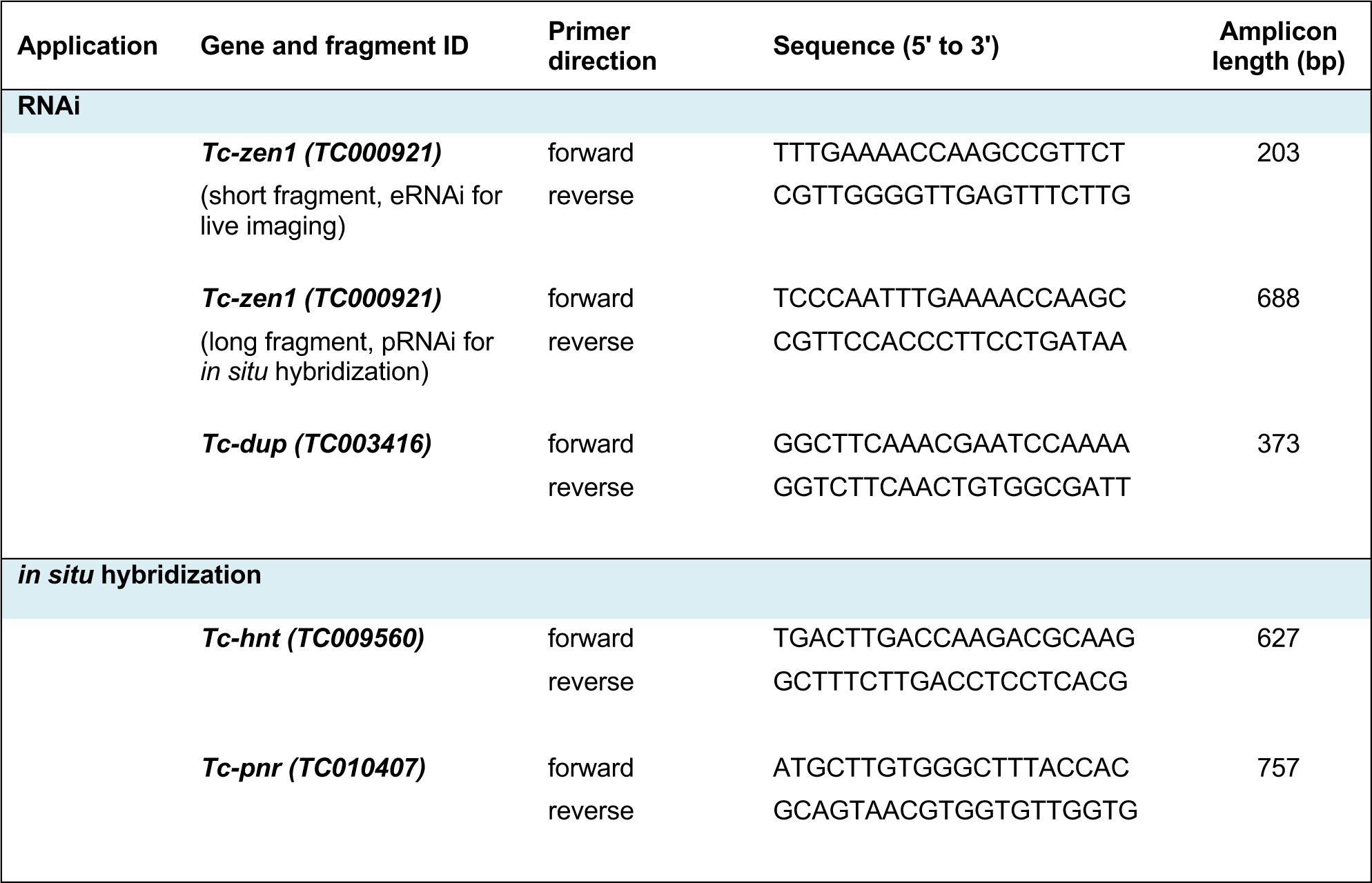
Primers used in this study.

## Notes

### Competing Interest Statement

The authors have declared no competing interest.

